# A facile immunopeptidomics workflow for capturing the HLA-I ligandome with PEAKS XPro

**DOI:** 10.1101/2021.05.20.444976

**Authors:** Kyle S. Hoffman, Baozhen Shan, Jonathan R. Krieger

## Abstract

Identifying antigens displayed specifically on tumour cell surfaces by human leukocyte antigen (HLA) proteins is important for the development of immunotherapies and cancer vaccines. The difficulty in capturing an HLA ligandome stems from the fact that many HLA ligands are derived from splicing events or contain mutations, hindering their identification in a standard database search. To address this challenge, we developed an immunopeptidomics workflow with PEAKS XPro that uses *de novo* sequencing to uncover such peptides and identifies mutations for neoantigen discovery. We demonstrate the utility of this workflow by re-analyzing HLA-I ligandome datasets and reveal a vast diversity in peptide sequences among clones derived from a colorectal cancer tumour. Over 8000 peptides predicted to bind HLA-I molecules were identified by *de novo* sequencing only (not found in the UniProt database) and make up over 50% of identified peptides from each sample. Lastly, tumour-specific mutations and consensus sequence motif characteristics are defined. This workflow is widely applicable to any immunopeptidomic mass spectrometry dataset and does not require custom database generation for neoantigen discovery.

## Introduction

The display of peptide antigens on cell surfaces by the major histocompatibility complex (MHC) plays an important role in adaptive immunity through recognition by CD8+ T cells. The antigens are produced from proteasome degradation of proteins and peptidase trimming into short peptides 8-12 amino acids in length (reviewed in (1, 2)). In the endoplasmic reticulum the antigens bind to human leukocyte antigen (HLA) molecules that make up the MHC, which is then exported to the cell surface. Here, the displayed antigens can be recognized by CD8+ T cells, thus initiating a cytolytic response that will eliminate the antigen-presenting cell.

Immunopeptidomics involves isolating MHC bound peptides by immunoaffinity purification followed by liquid chromatography tandem mass spectrometry (LC-MS/MS) for identification. A major challenge in this field is to identify novel antigens, or neoantigens, which are not directly encoded from genetic sequences and are absent from standard databases. A significant fraction (approximately 30%) of the antigens are *cis*- and *trans*-spliced peptides (3), or are non-canonical peptides derived from noncoding regions or frameshifted genes (4, 5). Identifying peptides derived from non-canonical reading frames typically requires a custom database to be generated from RNA-sequencing, followed by searching LC-MS/MS data against the custom database. Despite much success with this approach, RNA-sequencing adds significant time and cost, and *cis*- and *trans*-spliced peptides may not be identified in this manner.

To overcome these challenges, we have developed an immunopeptidomics workflow that identifies HLA ligands not found in the UniProt database and does not require next-generation sequencing. This aids in the discovery of neoantigens containing somatic mutations, peptides derived from splicing events, or non-canonical proteins. The workflow relies on *de novo* sequencing to retrieve peptide sequences directly from the MS data and is combined with a homology-based database search to identify missense mutations. Using highly stringent parameters in our MS analysis, and predicting HLA-I binding peptides with NetMHCpan-4.1, we improve HLA-I ligand identification with high confidence and characterize diverse sets of peptides across colorectal cancer tumour clones from a previously published dataset (6).

## Results

To demonstrate the utility of our immunopeptidomics workflow we re-analyzed the ligandome data from Demmers *et al*. (6) and demonstrate how it can capture the diversity in HLA-I ligands presented on different organoids from the same tumour. The genetic diversity among clones of the same tumour is an important driver of tumour evolution and acquiring resistance against chemotoxic stress. Tumour heterogeneity is commonly attributed to differential drug-resistance among tumour clones (7). For this reason, it is necessary to develop personalized multi-peptide vaccines to eliminate tumour clones that would otherwise evade immunotherapy. Immunopeptidomics is a valuable method for characterizing the antigens displayed on the surface of tumour cells by HLA-I molecules. As described in (6), by sampling different organoids from the same tumour of a patient and defining inter-clonal differences in HLA-I ligands, more effective immunotherapy drugs can be developed.

In the paper by Demmers *et al.,* four tumour cell clones are taken from the tumour of a colorectal cancer patient, along with a sample from normal colon tissue. Each isolated clone is amplified in conditioned medium to prevent cell differentiation and expanded into an organoid, thereby increasing the abundances of HLA-I ligands for more accurate and sensitive identification (6). After 8 weeks of organoid expansion the cells are harvested, peptide-HLA complexes are immunoprecipitated, and the bound peptides are isolated using size-exclusion filters. Each sample is then analyzed by LC-MS/MS.

In our immunopeptidomics workflow we use PEAKS XPro software for *de novo* sequencing, database searching, and for identifying mutations with the Spider algorithm. The peptides that pass a 1% FDR cutoff and *de novo* sequences with >80% average local confidence scores are kept so that only unique, high-quality peptides of 8-12 amino acids in length are considered. All mutations discovered in our analysis are supported by a mutation ion intensity >1%. In other words, a pair of major fragment ions (b- or y-ions) must be found before and after the amino acid with a minimum relative ion intensity of 1% in each spectrum to ensure confidence in the identity of the mutation. Modified forms of a peptide are also removed so that each peptide sequence is unique (Figure 1).

**Figure 1.**
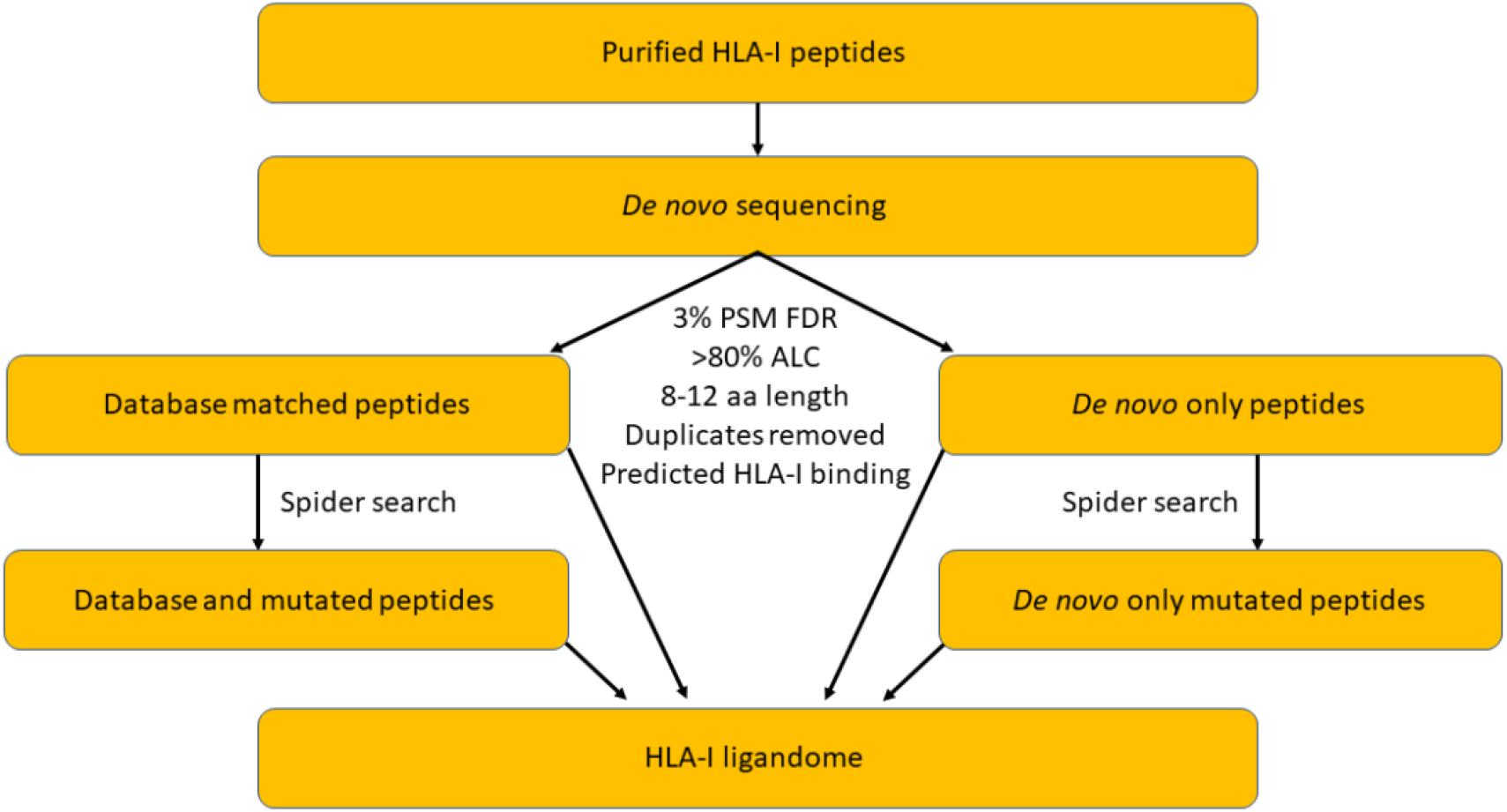
Flow chart of immunopeptidomics workflow with PEAKS XPro.

We then performed HLA-I ligand predictions using the NetMHCpan-4.1 server (8). This method uses artificial neural networks trained from eluted ligands and binding affinity data to accurately predict peptide binding to known HLA molecules (8). We chose to use this HLA-I peptide-binding prediction tool since it outperforms other machine learning-based tools (9). Peptides were assigned as HLA-I ligands if their eluted ligand score ranks within the top 2% out of a large set of random natural peptides. Our stringent peptide filters led to a high portion of assignment of predicted HLA-I ligands (parameters used for MS analysis are listed in Table 1). Approximately 90% of all database identified peptides and 70% *de novo* only sequenced peptides were predicted to bind one or more of the HLA-I proteins (Table 2). In each of the tumour clones and in the normal tissue sample, greater than 50% of predicted binders were identified by *de novo* sequencing only (Figure 2a), demonstrating that database searching alone is insufficient for capturing the full HLA-I ligand landscape.

**Table 1.** Parameters used for analysis of HLA-I peptides.

| Parameters | Settings |
| --- | --- |
| Variable PTMs | Oxidation (M), Cysteinylation |
| Mass error tolerance | 10 ppm precursor ion<br>0.02 Da fragment ion |
| Enzyme specificity | None |
| Fragmentation mode | HCD, EThcD |
| Database | Unreviewed Human Swiss/UniProt<br>(December 2<sup>nd</sup>, 2020) |
| Contaminant database | cRAP and FBS |
| De novo ALC% | >80% |
| Peptide length | 8-12 aa |
| Peptide/PSM FDR | 3%/1% |
| Duplicates | removed |

**Table 2.**
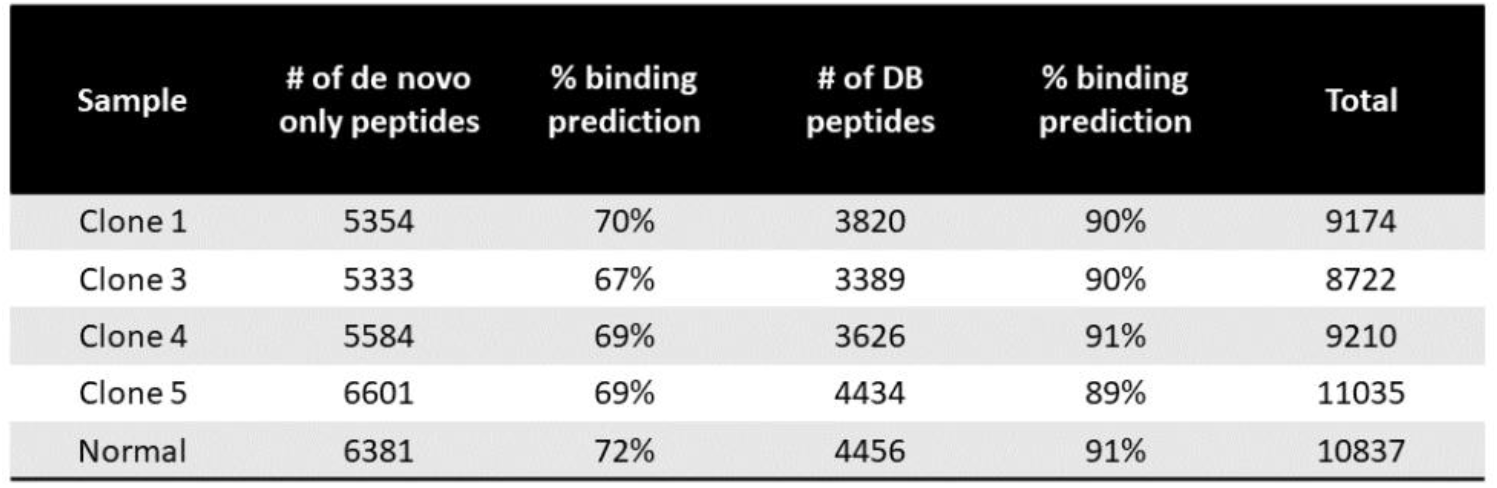
Percent binding predictions for peptides identified by *de novo* sequencing only or database searching.

| Sample | # of de novo only peptides | % binding prediction | # of DB peptides | % binding prediction | Total |
| --- | --- | --- | --- | --- | --- |
| Clone 1 | 5354 | 70% | 3820 | 90% | 9174 |
| Clone 3 | 5333 | 67% | 3389 | 90% | 8722 |
| Clone 4 | 5584 | 69% | 3626 | 91% | 9210 |
| Clone 5 | 6601 | 69% | 4434 | 89% | 11035 |
| Normal | 6381 | 72% | 4456 | 91% | 10837 |

**Figure 2.**
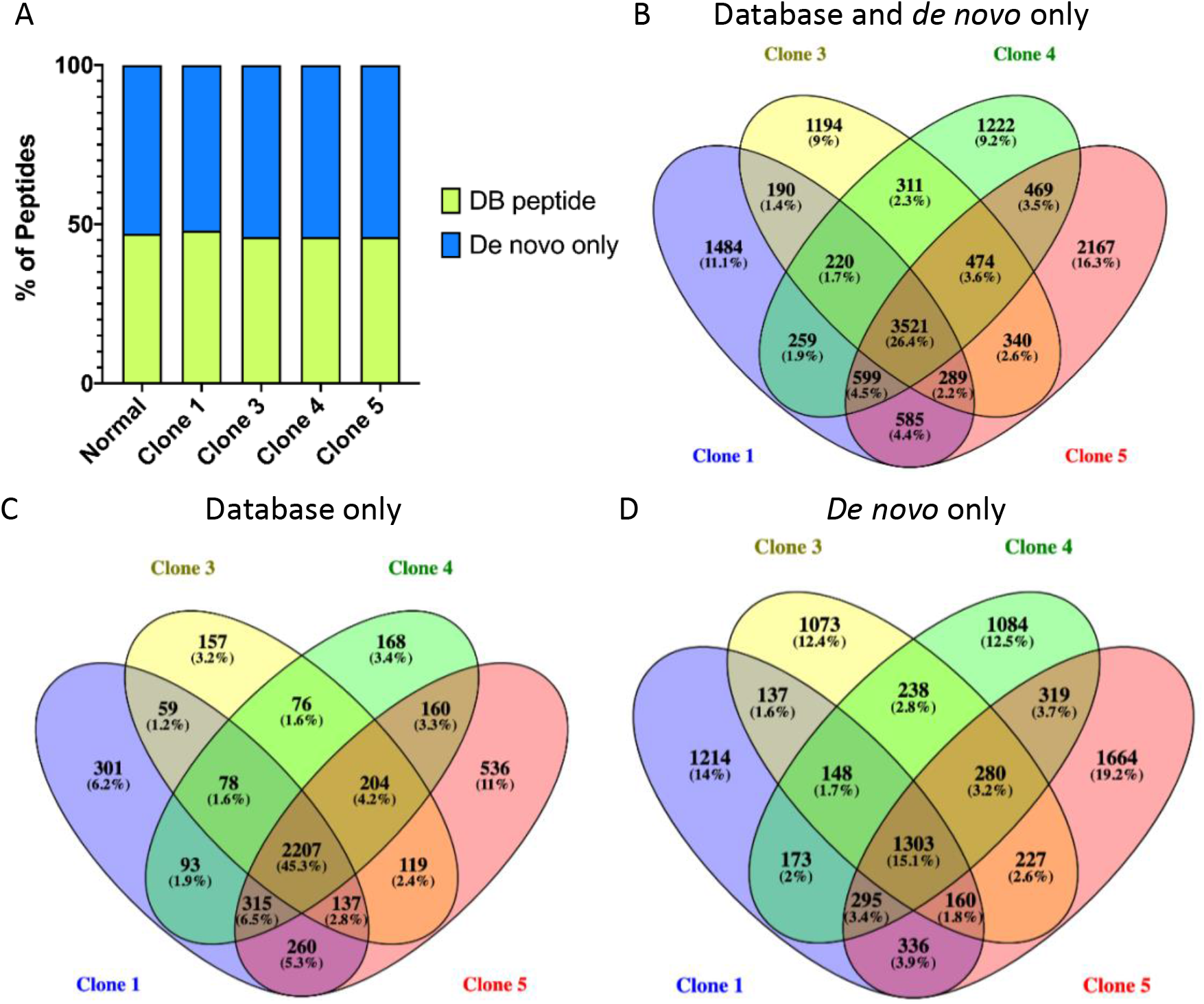
HLA-I predicted ligands identified in database searches or by *de novo* sequencing only. A) The percentage of peptides for each sample that were identified by a database search (green) or by de novo sequencing only (blue) and are predicted as HLA-I binders. B-D) Venn diagrams showing the diversity of HLA-I peptides identified by database searching and *de novo* sequencing (B), database searching alone (C), or by *de novo* sequencing only (D).

Next, we compared the predicted HLA-I binding peptides across tumour clones to determine the diversity of peptides that may be present among clones from the same colorectal cancer tumour. Among all peptides identified in tumour samples, 26.4% are common to all clones, whereas the peptides unique to any one clone make up between 9-16% of peptides (Figure 2b). When comparing database identified peptides, over 45% of all peptides are shared among tumour clones (Figure 2c). A greater diversity of peptides is found from *de novo* sequencing data. Only 15% of peptides are shared while over 1000 peptides are unique to any individual tumour organoid (Figure 2d). This outlines the importance of *de novo* sequencing in capturing HLA-I ligand diversity across different clones and suggests that many of the peptides, potentially derived from splicing events or non-canonical proteins, are absent from a standard database such as UniProt.

With the goal of identifying colorectal tumour-specific peptides for vaccine development, we compared the predicted HLA-I ligands identified in tumour clones to those present in the normal colon tissue sample. On average, 39% of peptides were common between each tumour clone and the normal sample, while 36% and 25% of peptides were specific to normal and tumour samples, respectively (Figure 3a). We next looked for differences in the length distribution between tumour-specific and normal peptides. The predominant peptide length was nine amino acids across all samples, which is characteristic of HLA-I binding peptides (Figure 3b). We noticed a slight increase in 10mer and 12mer peptides in tumour-specific HLA-I ligands compared to normal tissue ligands, consistent with what was observed in Demmers *et al*. (6).

**Figure 3.**
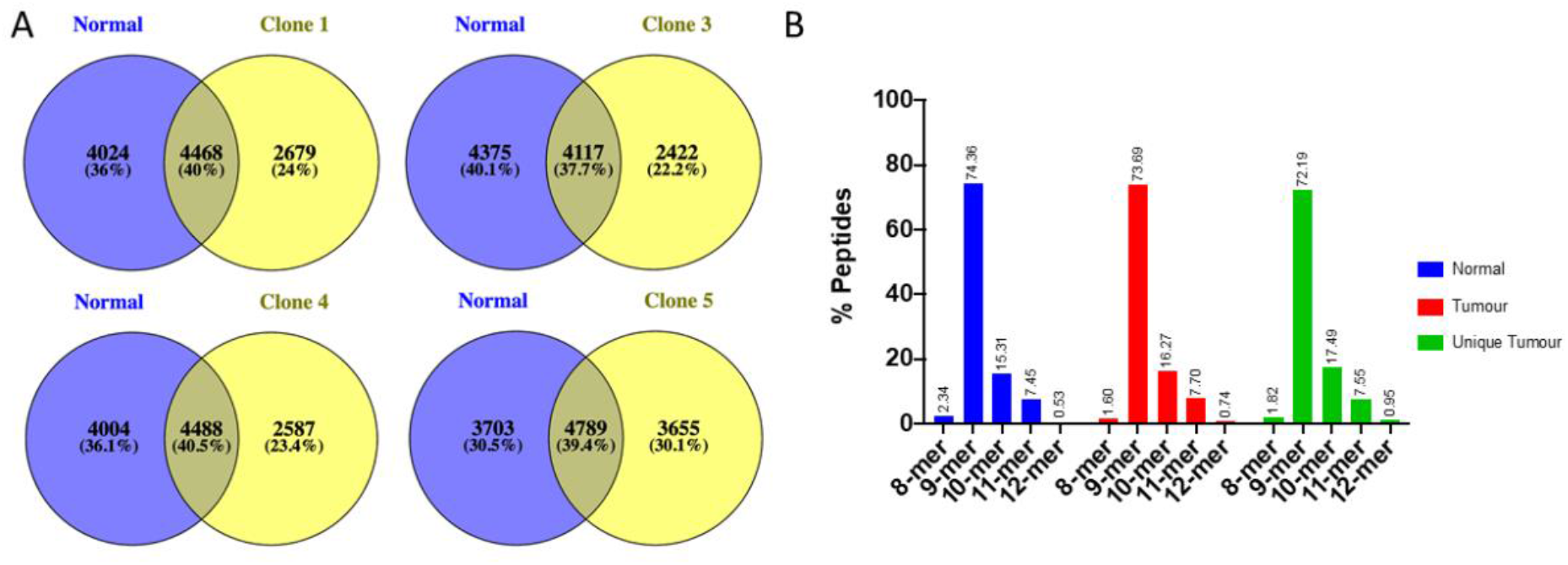
Characterization of HLA-I ligands from normal and tumour samples. A) All HLA-I peptides identified in each tumour sample were compared to HLA-I peptides from normal colorectal organoids. B) Length distribution of HLA-I peptides from normal organoid clones (blue), tumour clones (red), and those unique to tumour clones (green).

When generating sequence motifs with the MixMHCp 2.1 tool (10, 11) using peptides predicted to bind HLA proteins (HLA-A*02:01, HLA-B*15:01/57:01, HLA-C*03:04/06:02), tumour-specific amino acid preferences were observed (Figure 4). There were no significant differences in amino acid preferences in sequence motif positions generated for HLA-A*02:01, HLA-C*03:04, or HLA-C*06:02. Interestingly, there was a strong preference for tyrosine at the last position of the 9-mer peptide motif for HLA-B*15:01 and HLA-B*57:01 in the normal tissue sample. Tyrosine was not a common residue at the C-terminus of tumour-specific sequence motifs for these HLA-I alleles (Figure 4). Moreover, lysine and proline were most common at positions one and four in the HLA-B*57:01 sequence motif for the normal tissue samples, whereas in tumour samples leucine and valine occur more frequently at the first position, and the preference for proline was less pronounced. Taken together, tumour-specific preferences were observed for HLA-B sequence motifs which favour smaller hydrophobic residues at the N- and C-terminus over lysine and tyrosine, respectively, and a stronger preference for proline was found in normal tissue samples at position 4 in HLA-B*57:01 motif.

**Figure 4.**
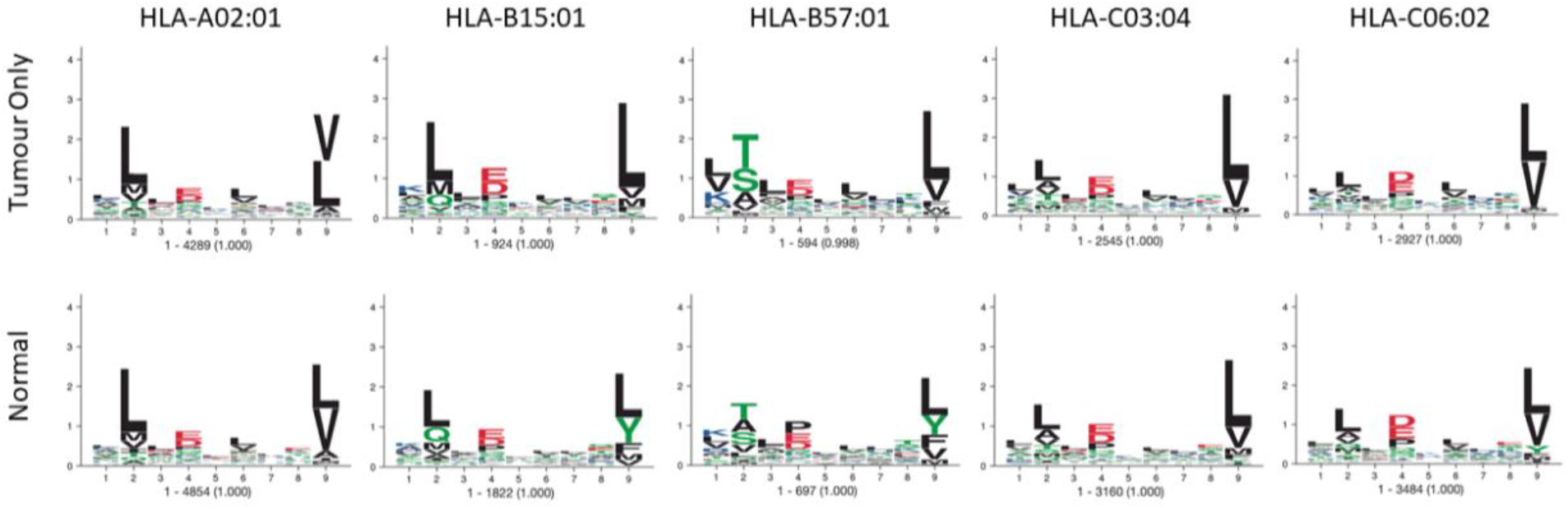
Sequence motifs for HLA-I allele from normal and tumour organoid clones. HLA-I peptides either tumour-specific or detected in normal organoid clones were used to generate sequence motifs for each of the patients’ HLA-I alleles using MixMHCp 2.1.

Lastly, we analyzed the data using PEAKS XPro Spider which allows for mismatches in a homology-based database search to identify missense mutations. Peptides containing cancer-associated somatic mutations and neoantigen discovery are of interest for the development of an immunotherapy that specifically targets cancer cells. We considered mutations that were supported by fragmentation ions with peak intensities greater than or equal to 1% in each peptide spectrum and found in peptides with lengths between 8 and 12 amino acids. Of the seven mutated peptides that met these criteria, two were identified in cancer clones only. One of the two peptides contained a Q1846D mutation in Laminin subunit alpha-3 but was not predicted to bind any of the expressed HLA-I proteins. The second was a peptide nine amino acids in length and contained a V29A mutation in cyclin-dependent kinase 3 (CDK3; Figure 5). The mutated peptide had an eluted ligand rank of 2.59% for HLA-A*02:01, slightly above the 2% threshold used for ligand binding predictions. Nonetheless, this mutation was not present in normal tissue samples and identified only in tumour clones three, four, and five. Moreover, CDK3 is highly expressed in colorectal cancer and plays a role in cancer cell proliferation and migration (12).

**Figure 5.**
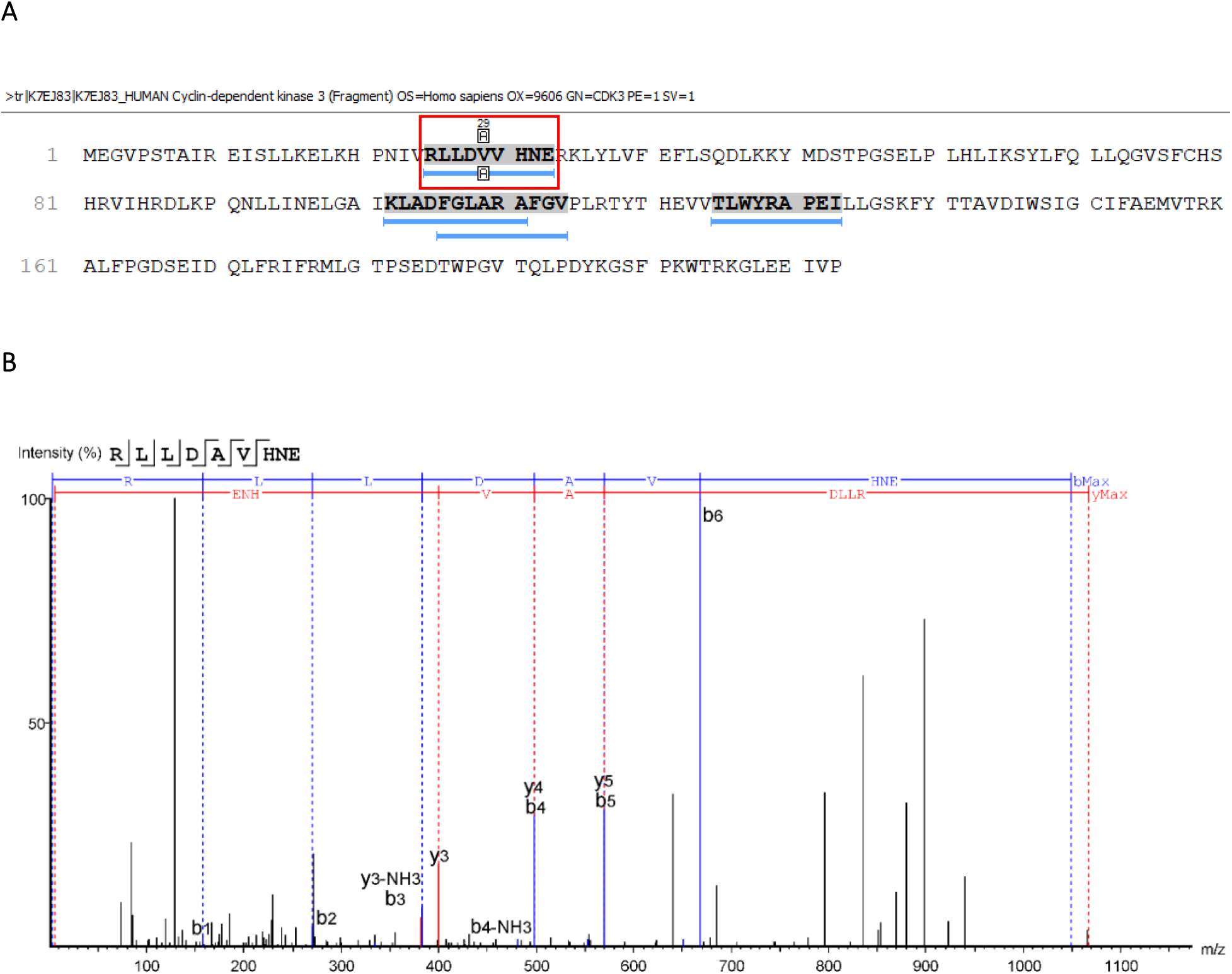
Mutation identified in CDK3 specific to colorectal cancer tumour-clones. A) Protein sequence of CDK3 showing the peptides mapped to the sequence in a database search. The mutated peptide is outlined in red, and the position of the mutation is shown above the sequence. B) A representative tandem mass spectrum supporting the V29A mutation in the peptide outlined in A).

## Discussion

In this study, we have developed an immunopeptidomics workflow with PEAKS XPro combined with NetMHCpan 4.1 to identify HLA-I ligands strictly from MS data. We demonstrate that *de novo* sequencing can reveal a diverse set of peptides which could not be discovered in a typical database search and is necessary for capturing HLA-I ligand diversity across organoid clones from the same tumour. We revealed differences between tumour and normal tissue samples in amino acid frequencies at specific positions within sequence motifs. In addition, a mutated peptide unique in tumour clones was identified in CDK3 using the PEAKS XPro Spider tool, which may represent a neoantigen identification. The tumour-specific amino acid preferences and CDK3 mutation will be important to consider in the development of an immunotherapy or vaccine against colorectal cancer tumours.

The immunopeptidomics workflow presented here was designed to be widely applicable and straightforward to use, while maintaining high accuracy and sensitivity in HLA ligand detection. The large diversity of peptide sequences identified across tumour clones was uncovered by *de novo* sequencing. While we acknowledge that these peptides cannot be accurately matched to sequences in a database to help define their biosynthetic routes (*cis*- or *trans*-splicing) or frameshift mutations, the workflow allows for a facile method for their identification which does not rely on custom database generation or RNA-sequencing. Future studies will focus on statistical significance methods to help identify *cis*- or *trans*-spliced peptides from the *de novo* only peptides that have partial matches to database sequences.

In conclusion, PEAKS XPro coupled with web-based tools such as NetMHCpan 4.1 (8) and mixMHCp (10, 11) provides a sensitive, accurate, and user-friendly workflow for the discovery of peptides displayed on the cell surface by MHC complexes. This workflow can be applied to any immunopeptidomic MS dataset to improve peptide identification, HLA ligand prediction, and potentially discover neoantigens for the development of immunotherapies.

## Methods

### LC-MS/MS Analysis

All peptidomics raw files were downloaded from the ProteomeXchange consortium under the data set identifier PXD016582. The raw files were processed and re-analyzed using PEAKS Studio XPro with instrument parameters set to Orbitrap, HCD or EThcD fragmentation, and DDA acquisition mode. Associate feature with chimera scan was enabled to detect co-eluting peptides present within the same MS scan. Error tolerance was set to 10.0 ppm for precursor ions and 0.02 Da for fragmentation ions. Digestion was set as unspecific with no enzyme selection. Oxidation of methionine and cysteinylation were set as variable modifications. The data were searched against the unreviewed human UniProt database (December 2, 2020), as well as the cRAP and cRFP (13) databases to remove contaminating peptides. False discovery rate (FDR) was set to 3% for peptide identification (or 1% PSM FDR) and estimated using the decoy-fusion method (14). *De novo* average local confidence score was set to greater than 80%. Only peptides equal to or between lengths of 8-12 amino acids were included in the analysis.

### Analysis of HLA-I Peptides

NetMHCpan 4.1 (8) was used to predict the binding affinity for HLA-I peptides. The colorectal cancer patients’ HLA-I alleles were obtained from Demmers *et al*. (6). Peptides with an eluted ligand score of less than or equal to 2% were considered to bind HLA-I molecules. MixMHCp 2.1 (10, 11) was used to generate sequence motifs with a core length of nine amino acids using the peptides predicted to bind each of the HLA-I molecules. Venny 2.1 was used to create all Venn diagrams (15). Prism Version 9.0.2 was used to generate bar graphs.

## Acknowledgements

This work was supported by Bioinformatics Solutions Inc. We thank Amber Park for reviewing the manuscript.

## Author contributions

Conceptualization, K.S.H., J.R.K., B.S.; Writing, K.S.H.; Review and editing, J.R.K., B.S.

## Notes

### Competing Interest Statement

KSH, JRK, BS are employees of Bioinformatics Solutions Inc.

## References

1. Neefjes, J., Jongsma, M.L.M., Paul, P. and Bakke, O. (2011) Towards a systems understanding of MHC class i and MHC class II antigen presentation. Nat. Rev. Immunol., 10.1038/nri3084.

2. Groettrup, M., Kirk, C.J. and Basler, M. (2010) Proteasomes in immune cells: More than peptide producers? Nat. Rev. Immunol., 10.1038/nri2687.

3. Liepe, J., Marino, F., Sidney, J., Jeko, A., Bunting, D.E., Sette, A., Kloetzel, P.M., Stumpf, M.P.H., Heck, A.J.R. and Mishto, M. (2016) A large fraction of HLA class I ligands are proteasome-generated spliced peptides. Science (80-.)., 10.1126/science.aaf4384.

4. Laumont, C.M., Daouda, T., Laverdure, J.P., Bonneil, É., Caron-Lizotte, O., Hardy, M.P., Granados, D.P., Durette, C., Lemieux, S., Thibault, P., et al. (2016) Global proteogenomic analysis of human MHC class I-associated peptides derived from non-canonical reading frames. Nat. Commun., 10.1038/ncomms10238.

5. Ruiz Cuevas, M.V., Hardy, M.-P., Holly, J., Bonneil, É., Durette, C., Courcelles, M., Lanoix, J., Côté, C., Staudt, L.M., Lemieux, S., et al. (2021) Most Non-Canonical Proteins Uniquely Populate the Proteome or Immunopeptidome. Cell Rep.,https://doi.org/10.1016/j.celrep.2021.108815.

6. Demmers, L.C., Kretzschmar, K., Van Hoeck, A., Bar-Epraïm, Y.E., van den Toorn, H.W.P., Koomen, M., van Son, G., van Gorp, J., Pronk, A., Smakman, N., et al. (2020) Single-cell derived tumor organoids display diversity in HLA class I peptide presentation. Nat. Commun., 10.1038/s41467-020-19142-9.

7. Dagogo-Jack, I. and Shaw, A.T. (2018) Tumour heterogeneity and resistance to cancer therapies. Nat. Rev. Clin. Oncol., 10.1038/nrclinonc.2017.166.

8. Reynisson, B., Alvarez, B., Paul, S., Peters, B. and Nielsen, M. (2020) NetMHCpan-4.1 and NetMHCIIpan-4.0: improved predictions of MHC antigen presentation by concurrent motif deconvolution and integration of MS MHC eluted ligand data. Nucleic Acids Res., 10.1093/nar/gkaa379.

9. Mei, S., Li, F., Leier, A., Marquez-Lago, T.T., Giam, K., Croft, N.P., Akutsu, T., Ian Smith, A., Li, J., Rossjohn, J., et al. (2020) A comprehensive review and performance evaluation of bioinformatics tools for HLA class I peptide-binding prediction. Brief. Bioinform., 10.1093/bib/bbz051.

10. Bassani-Sternberg, M. and Gfeller, D. (2016) Unsupervised HLA Peptidome Deconvolution Improves Ligand Prediction Accuracy and Predicts Cooperative Effects in Peptide–HLA Interactions. J. Immunol., 10.4049/jimmunol.1600808.

11. Gfeller, D., Guillaume, P., Michaux, J., Pak, H.-S., Daniel, R.T., Racle, J., Coukos, G. and Bassani-Sternberg, M. (2018) The Length Distribution and Multiple Specificity of Naturally Presented HLA-I Ligands. J. Immunol., 10.4049/jimmunol.1800914.

12. Yang, C. and Sun, J.J. (2015) Mechanistic studies of cyclin-dependent kinase inhibitor 3 (CDKN3) in colorectal cancer. Asian Pacific J. Cancer Prev., 10.7314/APJCP.2015.16.3.965.

13. Shin, J., Kwon, Y., Lee, S., Na, S., Hong, E.Y., Ju, S., Jung, H.G., Kaushal, P., Shin, S., Back, J.H., et al. (2019) Common Repository of FBS Proteins (cRFP) to Be Added to a Search Database for Mass Spectrometric Analysis of Cell Secretome. J. Proteome Res.,10.1021/acs.jproteome.9b00475.

14. Zhang, J., Xin, L., Shan, B., Chen, W., Xie, M., Yuen, D., Zhang, W., Zhang, Z., Lajoie, G.A. and Ma, B. (2012) PEAKS DB: De novo sequencing assisted database search for sensitive and accurate peptide identification. Mol. Cell. Proteomics, 11, 1–8.

15. Oliveros, J.C. (2007) VENNY. An interactive tool for comparing lists with Venn Diagrams. http://bioinfogp.cnb.csic.es/tools/venny/index.html. bioinfogp.cnb.csic.es/tools/venny/index.html.

